# The cytidine deaminase APOBEC3G drives cancer mutagenesis and clonal evolution in bladder cancer

**DOI:** 10.1101/2022.09.05.503899

**Authors:** Weisi Liu, Kevin P. Newhall, Francesca Khani, LaMont Barlow, Duy Nguyen, Lilly Gu, Ken Eng, Bhavneet Bhinder, Manik Uppal, Charlotte Récapet, Andrea Sboner, Susan R. Ross, Olivier Elemento, Linda Chelico, Bishoy M. Faltas

## Abstract

Mutagenic processes leave distinct signatures in cancer genomes. The mutational signatures attributed to APOBEC3 cytidine deaminases are pervasive in human cancers. However, data linking individual APOBEC3 proteins to cancer mutagenesis *in vivo* are limited. Here, we show that transgenic expression of human APOBEC3G promotes mutagenesis, genomic instability, and kataegis, leading to shorter survival in a murine bladder cancer model. Acting as mutagenic fuel, APOBEC3G increases the clonal diversity of bladder cancers, driving divergent cancer evolution. We characterize the single base substitution signature induced by APOBEC3G *in vivo*, showing the induction of a mutational signature different from that caused by APOBEC3A and APOBEC3B. Analysis of thousands of human cancers reveals the contribution of APOBEC3G to the mutational profiles of multiple cancer types, including bladder cancer. Our findings define the role of APOBEC3G in cancer mutagenesis and clonal heterogeneity. These results potentially inform future therapeutic efforts that restrict tumor evolution.

## Introduction

Cancer evolution, heterogeneity, and treatment resistance are often linked to the acquisition of somatic mutations and chromosomal instability through various mutagenic processes that leave distinct signatures within cancer genomes(1). Somatic mutational signatures enriched with cytidine to thymine or guanine substitutions in TCW motifs (W: A or T) are prevalent in the genomes of several human cancers, particularly urothelial bladder cancer(1–5). These signatures have been associated with the apolipoprotein B mRNA-editing enzyme catalytic subunit 3 (APOBEC3) family(6,7), which encodes seven APOBEC3 proteins (APOBEC3A, B, C, D, F, G, H) in humans(8,9). APOBEC3 proteins have overlapping biochemical activity(10), confounding cause-and-effect experiments aiming to dissect the mutagenic impact of individual APOBEC3 proteins in human cells. However, there is only limited experimental data linking individual APOBEC3 proteins to known mutational signatures in animal cancer models(11). Consequently, the role of individual APOBEC3 proteins in driving mutagenesis, cancer cell fitness, and clonal evolution *in vivo* remains unclear. These knowledge gaps have impeded efforts to therapeutically target mutagenic processes to alter the fitness of cancer cells(12–14).

APOBEC3G is a double deaminase domain protein that is ubiquitously expressed in normal and cancer cells(15). Its canonical function is to restrict retroviruses(16–19), but its role in mutating genomic DNA in cancer cells has been unknown prior to our study. In this study, we show that transgenic human APOBEC3G expression significantly promotes genomic instability by increasing tumor mutational burden and copy-number alterations in a bladder cancer mouse model. These APOBEC3G-induced genomic lesions act as ’mutagenic fuel’ to drive clonal divergence and intratumor heterogeneity. We characterize a single base substitution signature induced by APOBEC3G *in vivo* and describe its contribution to the mutational profiles of thousands of human cancers.

## Results

### APOBEC3G transgene expression in a murine bladder cancer model

In contrast to humans, mice have a single *Apobec3* (*mA3*) gene(10). Therefore, knocking out the mouse *Apobec3* provides a null background to study the effects of each of the seven individual transgenic human APOBEC3 proteins. We used mice that constitutively express the human *APOBEC3G* (*hA3G*) transgene in the *Apobec3*-null background (*hA3G*(+) *mA3*(−/−)) to dissect the mutagenic role of APOBEC3G *in vivo* (**Fig. 1A**)(20–22) (**Methods**). In the hA3G(+) mA3(−/−) mice, transgenic APOBEC3G is expressed in multiple tissues, including the lung, the spleen, and immune cells(21,22). To confirm the expression of transgenic APOBEC3G in urinary bladders, we examined APOBEC3G mRNA level in the urinary bladder, which was comparable to the APOBEC3G mRNA levels in the lung and spleen (**Fig. 1B**). The protein expression of APOBEC3G in the urinary bladder was confirmed by SDS-PAGE western blotting (**Fig. 1C**). We reasoned that APOBEC3-induced mutagenesis alone would not be sufficient for tumor initiation(3,23,24), so we used the chemical carcinogen N-butyl-N-(4-hydroxybutyl)nitrosamine (BBN) to initiate tumorigenesis. We then compared the survival outcomes between the *hA3G*(+) *mA3*(−/−) mice and their *mA3*(−/−) littermate controls (**Fig. 1A**). In the 44 male mice enrolled in the experiment, blinded histopathologic examination showed that 19/25 (76%) *hA3G*(+) *mA3*(−/−) mice and 10/19 (52.6%) of *mA3*(−/−) mice developed bladder cancer (**Fig. 1D and 1E**). Strikingly, the *hA3G*(+) *mA3*(−/−) mice had significantly shorter survival compared to the *mA3*(−/−) mice (log-rank *P*=0.007) (**Fig. 1F; Supplementary Table S1**). We also included a cohort of C57BL/6J mice that were exposed to BBN as controls. The C57BL/6J mice, which harbor wild-type mouse *Apobec3*, developed a higher percentage of advanced bladder cancers and had significantly lower survival than the *mA3*(−/−) mice (log-rank *P*=0.0008) but phenocopied the *hA3G*(+) *mA3*(−/−) mice (**Supplementary Fig. S1**). All expired mice had bladder cancer.

**Figure 1.**
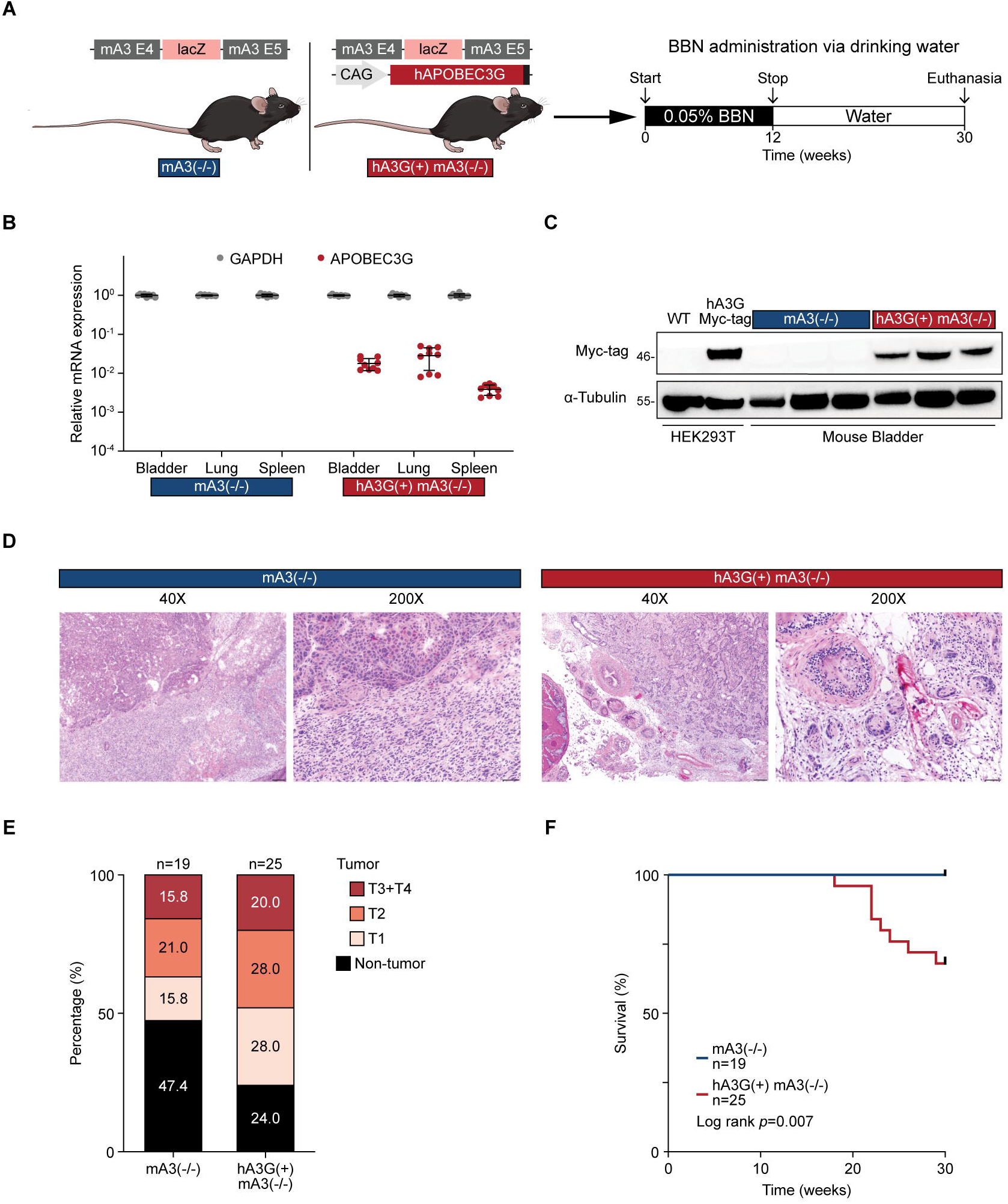
APOBEC3G contributes to carcinogenesis in a murine bladder cancer model. A) Experiment schema. The *hA3G*(+) *mA3(*−/−) mouse harbors a human *APOBEC3G* transgene under the control of a chicken beta-Actin promoter (CAG) on a *mA3*(−/−) background. The *mA3*(−/−) mouse was generated by knocking lacZ into the mAPOBEC3 locus between exome 4 and exome 5, resulting in the knockout of the mouse *Apobec3* gene. B) *APOBEC3G* mRNA is expressed in different organs (lung, bladder, and spleen) from the *hA3G*(+) *mA3*(−/−) mice but not the *mA3*(−/−) mice. Dots represent replicates from three different mice for each genotype. Horizontal lines indicate the mean value of mRNA expression. Error bars represent the mean ± SD. C) SDS-PAGE western blot. APOBEC3G protein is expressed in the bladders of the *hA3G*(+) *mA3*(−/−) mice but not in the *mA3*(−/−) mice. Exogenous expression of APOBEC3G in HEK293T cells is used as a control. D) Representative H&E images (40X and 200X) of bladder tumors in the *hA3G*(+) *mA3*(−/−) and *mA3*(−/−) mice. Scale bars represent 200μm in the 40X and 50μm in 200X images, respectively. E) Composite bar chart representing the percentage of tumor stages in mice from the *hA3G*(+) *mA3*(−/−) and *mA3*(−/−) groups. The non-tumor category includes benign tissue, hyperplasia, and dysplasia. F) Mice in the *hA3G*(+) *mA3*(−/−) (red) group had lower survival than mice in the *mA3*(−/−) (blue) group. Log-rank test. All dead mice had pathologically confirmed bladder cancers. *hA3G*: human *APOBEC3G*, *mA3*: mouse *Apobec3*.

### APOBEC3G drives genomic instability and clonal heterogeneity

To define the impact of human APOBEC3G on cancer mutagenesis, we performed deep whole-exome sequencing of somatic DNA from bladder cancers (mean 348x) and matched germline DNA from both the *hA3G*(+) *mA3*(−/−) and *mA3*(−/−) mice. The *hA3G*(+) *mA3*(−/−) tumors harbored a significantly higher mutational burden (median 95.7 mutations per Mb) than the *mA3*(−/−) tumors (median 48.5 mutations per Mb, *P*=0.04) (**Fig. 2A; Supplementary Table S2**). Because nuclear access is a prerequisite for APOBEC3G-induced mutagenesis, we examined the nuclear localization of APOBEC3G in a bladder cancer organoid established from a *hA3G*(+) *mA3*(−/−) tumor. SDS-PAGE western blotting showed that APOBEC3G was consistently present in the nuclear fraction (**Supplementary Fig. S2A**). The presence of APOBEC3G in the nuclear fraction was also confirmed in two human bladder cancer cell lines (5637 and RT112) using inducible GFP-tagged APOBEC3G (**Supplementary Fig. S2B and S2C**). In addition, to avoid obscuring the nuclear signal by the surrounding cytoplasmic APOBEC3G, we performed fluorescent imaging of GFP-tagged APOBEC3G within extracted RT112 nuclei (**Methods**) (**Supplementary Fig. S3A**). This confirmed the nuclear presence of APOBEC3G (**Supplementary Fig. S3B**). Finally, we performed fluorescent imaging and three-dimensional reconstruction of the extracted nuclei using confocal laser scanning microscopy (**Supplementary Fig. S3C and S3D; Supplementary Video S1**), demonstrating the presence of APOBEC3G in the nuclear compartment.

**Figure 2.**
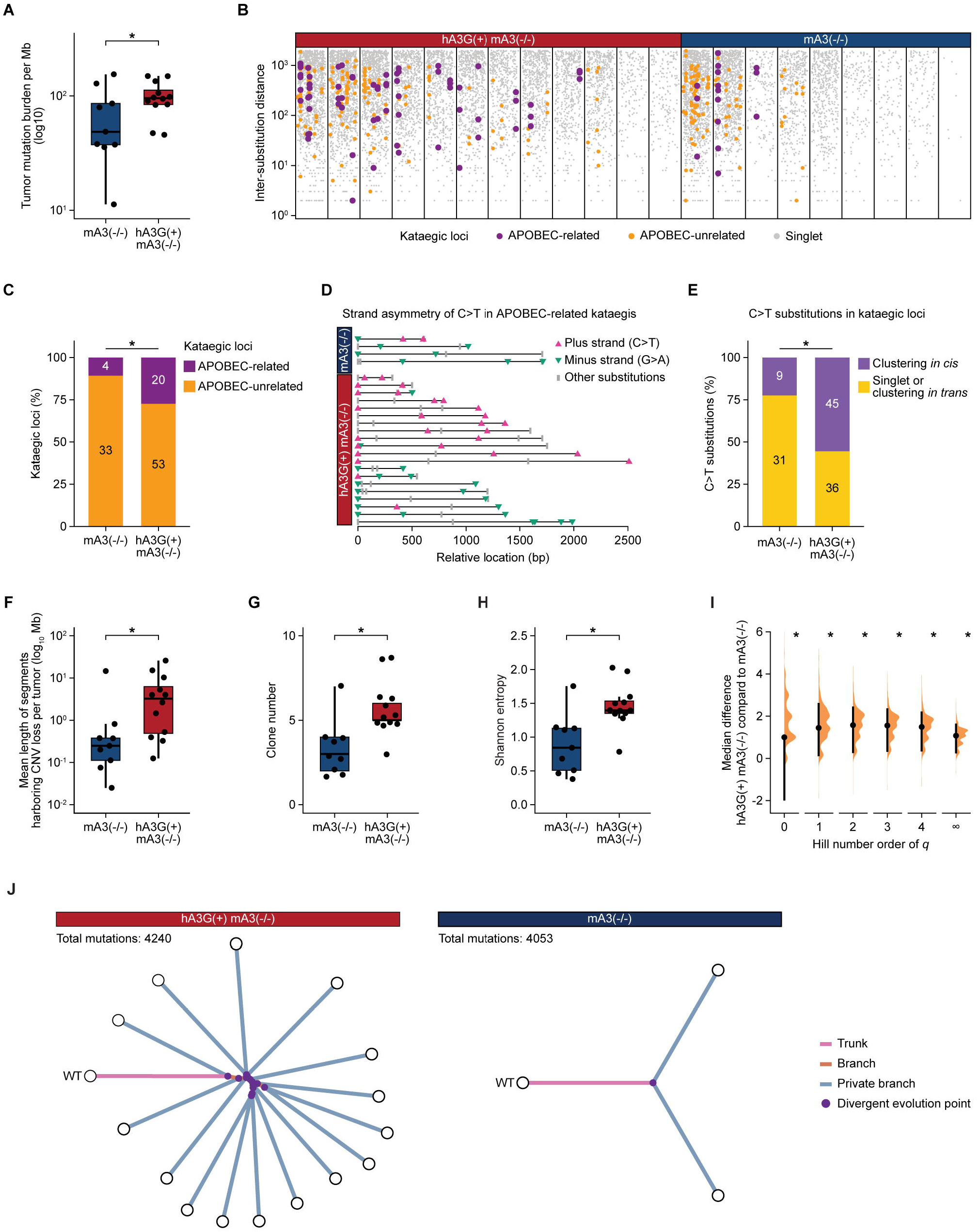
APOBEC3G increases genomic instability and intra-tumoral clonal heterogeneity. A) The *hA3G*(+) *mA3*(−/−) tumors harbor a higher mutational burden compared to the *mA3*(−/−) tumors. Mann-Whitney U test. *: *P*<0.05. The data is shown in the box plot as median with IQR. The lower whisker indicates Q1-1.5*IQR. The upper whisker indicates Q3+1.5*IQR. Each dot represents one tumor. B) Rainfall plot of kataegic loci in the *hA3G*(+) *mA3*(−/−) and *mA3*(−/−) tumors. Vertical lines represent individual tumors. Grey dots represent singlet substitutions. Orange dots indicate the substitutions within the APOBEC-unrelated kataegic loci. Purple dots indicate substitutions within the APOBEC-related kataegic loci with a significant number of C>T or G>A in kataegic loci calculated by binormal test (**Methods**). C) Bar chart representing the number of kataegic loci indicating that APOBEC-related kataegic loci were enriched in *hA3G*(+) *mA3*(−/−) tumors. Fisher’s exact test. *: *P*<0.05. D) Strand asymmetry of C>T substitutions in APOBEC-related kataegis. Each line represented an APOBEC-related kataegic loci in different genotypes. The length of the line indicates the relative distance between substitutions. The triangles with different directions and colors indicated the strandedness of C>T substitutions. E) Bar chart representing the number of C>T substitutions occurring *in cis* and the C>T substitutions occurring *in trans* and singlet in kataegic loci, indicating the C>T substitutions occurring *in cis* were enriched in *hA3G*(+) *mA3*(−/−) tumors. Fisher’s exact test. *: *P*<0.05. F) The *hA3G*(+) *mA3*(−/−) tumors had a broader mean length of CNV loss events compared to the *mA3*(−/−) tumors. Mann-Whitney U test. *: *P*<0.05. The data is shown in the box plot as the median with IQR. The lower whisker indicates Q1-1.5*IQR. The upper whisker indicates Q3+1.5*IQR. Each dot represents one tumor. G) The *hA3G*(+) *mA3*(−/−) tumors harbor a higher number of clones compared to the *mA3*(−/−) tumors. Mann-Whitney U test. *: *P*<0.05. Boxplots show the median and IQR. The lower whisker indicates Q1-1.5*IQR. The upper whisker indicates Q3+1.5*IQR. Individual dots indicate individual tumors. H) The *hA3G*(+) *mA3*(−/−) tumors displayed higher Shannon entropy compared to the *mA3*(−/−) tumors. Mann-Whitney U test. *: *P*<0.05. Boxplots show the median and IQR. The lower whisker indicates Q1-1.5*IQR. The upper whisker indicates Q3+1.5*IQR. Individual dots indicate individual tumors. I) The median difference in Hill number order between *hA3G*(+) *mA3*(−/−) and *mA3*(−/−) tumors are shown by the Gardner-Altman estimation plot. The mean difference is plotted as a bootstrap sampling distribution and depicted as a dot with a 90% confidence interval indicated by the ends of the vertical error bar. J) Phylogenetic trees for representative one *hA3G*(+) *mA3*(−/−) and one *mA3*(−/−) tumor with comparable total mutational burdens but divergent clonal evolution patterns. ‘WT’ represents the inferred wild-type genome. Branch length corresponds to the proportion of the number of shared variants. The length of the branches in different tumors is normalized to the same scale. CNV: copy number variant. IQR: interquartile range. *hA3G*: human *APOBEC3G*, *mA3*: mouse *Apobec3*.

We then investigated the presence of kataegis, a clustered mutagenesis process attributed to the processive deaminase activity of APOBEC3 proteins(5,6,25–27), including APOBEC3G(6). We hypothesized that human APOBEC3G deaminates proximate cytosines to generate kataegic loci in mouse tumors. We identified these kataegic loci based on the inter-substitution distance and the number of substitutions involved in each locus and defined the APOBEC-related kataegic loci based on strand coordination (**Methods**). We also used the computational tool SeqKat(28) to identify the kataegic loci (**Supplementary Fig. S4**). The *hA3G*(+) *mA3*(−/−) tumors harbored more APOBEC-related kataegis than the *mA3*(−/−) tumors (**Fig. 2B-D**). Moreover, the strand-coordinated C>T mutations occurring *in cis* in kataegic loci were enriched in *hA3G*(+) *mA3*(−/−) tumors compared to *mA3*(−/−) tumors (**Fig. 2E**). Collectively, these results indicate that APOBEC3G increases kataegis *in vivo*. We also examined APOBEC3-induced copy number changes using CNVkit(29) (**Methods**). The mean length of the copy number loss events (log2 copy ratio < −0.2) was longer in the *hA3G*(+) *mA3*(−/−) tumors compared to the *mA3*(−/−) tumors (*P*=0.007) (**Fig. 2F; Supplementary Fig. S5 and S6**).

Since mutational and structural genomic instability is a crucial driver of intra-tumoral heterogeneity and clonal evolution(30), we hypothesized that human APOBEC3G-induced mutations and copy number variants engender intra-tumoral heterogeneity in our murine bladder cancer model. First, we quantified the number and size of cancer clones in each tumor using two computational methods EXPANDS (31,32) and Pyclone-vi(33) (**Methods**). The *hA3G*(+) *mA3*(−/−) tumors harbored a significantly higher clone number compared to the *mA3*(−/−) tumors (Median 5 versus 3 clones, respectively, *P*=0.001) using Pyclone-vi (**Fig. 2G**). We then used Shannon entropy to represent phylogenetic diversity(34). Shannon entropy was significantly higher in the *hA3G*(+) *mA3*(−/−) tumors compared to the *mA3*(−/−) tumors (Median 1.3 versus 0.8, respectively, *P*=0.001), which indicated higher diversity in *hA3G*(+) *mA3*(−/−) tumors (**Fig. 2H**). Next, we compared the diversity profiles based on Hill numbers(34,35), which include a sensitivity parameter that controls the weighing of clones in each tumor to ensure that rare clones do not skew the comparisons **(Fig. 2I)**. This analysis confirmed that the *hA3G*(+) *mA3*(−/−) tumors harbor significantly higher clonal diversity compared to the *mA3*(−/−) tumors (**Fig. 2G and 2H**). These results were also confirmed by EXPANDS(31,32) (**Fig. 2J; Supplementary Fig. S7**). In summary, our findings suggest that human APOBEC3G contributes to intra-tumoral clonal heterogeneity.

### The mutational signature induced by APOBEC3G

The *hA3G*(+) *mA3*(−/−) tumors harbored a significantly higher number of C>T substitutions in the CC motif compared to the *mA3*(−/−) tumors (*P*=0.002) (**Fig. 3A**). This is consistent with the known CC motif preference of APOBEC3G-induced mutations in viral genomes(18,36–39). To further characterize the APOBEC3G-induced mutational signature *in vivo*, we deconstructed the trinucleotide mutational spectra for mutations in each tumor (**Supplementary Table S3**). C>T substitutions in the TCC, GCC, CCC, CCT, and GCG motifs were significantly higher in the *hA3G*(+) *mA3*(−/−) tumors than in the *mA3*(−/−) tumors. In contrast, C>T substitutions in the ACA motif significantly decreased (**Supplementary Fig. S8A**). In addition, APOBEC3G preferred the transcribed and lagging strands (**Supplementary Fig. S8B**). The preference for the lagging strand is similar to previous reports of mutagenesis by other APOBEC3 family members, including APOBEC3A and APOBEC3B(40,41).

**Figure 3.**
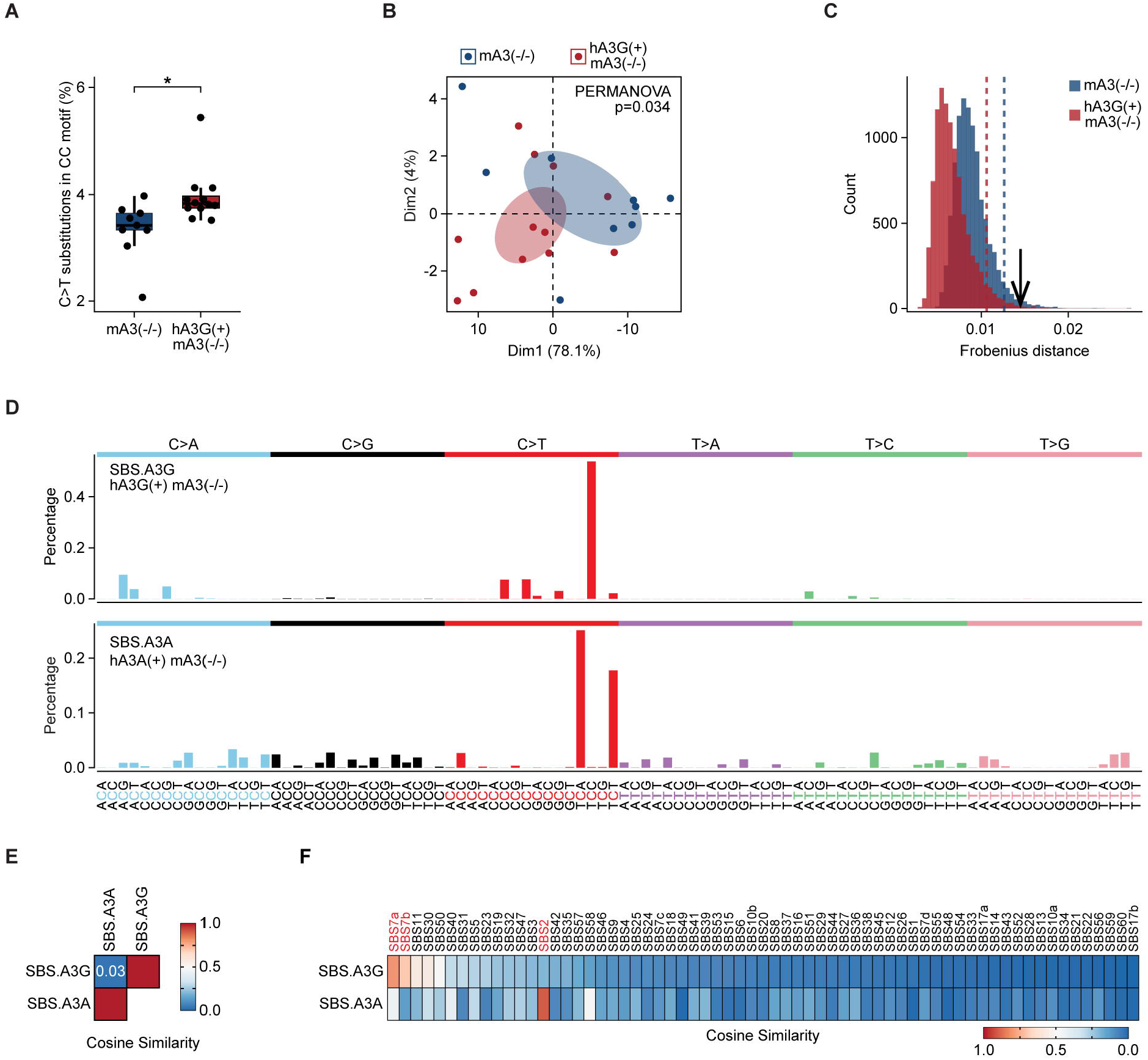
APOBEC3G generates a distinct *in vivo* mutational signature. A) The *hA3G*(+) *mA3*(−/−) tumors had higher C>T substitutions in the CC motif compared to the *mA3*(−/−) tumors. Mann-Whitney U test. *: *P*<0.05. Bars represent the mean percentage of C>T substitutions in the CC motif to total substitutions. Error bars represent SD. Each dot represents one tumor. B) PCA analysis based on the trinucleotide mutational spectrum showing divergence between the *hA3G*(+) *mA3*(−/−) and *mA3*(−/−) tumors. The confidence ellipses presented the 95% CI. Permutational Multivariate Analysis of Variance test was used to calculate the *P* value. C) Significant mutational shift (black arrow) between the centroids of bootstrapped mutational spectra of the *hA3G*(+) *mA3*(−/−) and *mA3*(−/−) tumors. The histogram represents the distribution of the difference between centroids of bootstrapped mutational spectra and original mutational spectra of the *hA3G*(+) *mA3*(−/−) tumors (red) and *mA3*(−/−) (blue). The dashed lines indicate the threshold of significance (*P*=0.05) for each genotype. D) The single base substitution signature induced by transgenic expression of human APOBEC3G (SBS.A3G) and transgenic expression of human APOBEC3A (SBS.A3A). E) SBS.A3G has a low cosine similarity to SBS.A3A. F) Cosine similarity of experimentally derived and COSMIC PCAWG single base substitution signatures. SBS.A3G has a low cosine similarity to SBS2 and SBS13. SBS.A3A had a high cosine similarity with SBS2. *hA3G*: human *APOBEC3G*, *mA3*: mouse *Apobec3*, CI: confidence interval. SBS: single base substitution, COSMIC: Catalogue Of Somatic Mutations in Cancer, PCAWG: the Pan-Cancer Analysis of Whole Genomes.

To characterize the differences in the mutational landscape between the tumors from the two genotypes, we performed PCA analysis of the 96 trinucleotide mutational spectra. The mutational spectra from *hA3G*(+) *mA3*(−/−) tumors showed significant divergence from *mA3*(−/−) tumors (*P*=0.03) (**Fig. 3B**). To confirm these results, we adapted a statistical framework(12,13) that employs bootstrap resampling of the 96 trinucleotide mutational profiles of sequenced tumors to extract the net mutational signature generated by human APOBEC3G *in vivo* (**Methods**). We found a significant mutational spectrum shift between tumors in the *hA3G*(+) *mA3*(−/−) and the *mA3*(−/−) groups (*P*=0.05) (**Fig. 3C**). We extracted the net signature of APOBEC3G (SBS.A3G) using the mutational shift distance (MSD) method (**Methods**). SBS.A3G was characterized by C>T substitutions in the TCC, CCC, and CCT (**Fig. 3D**). The TCC motif is consistent with the secondary motif preference of APOBEC3G for the TC context observed in the HIV genome(18,36,42,43) (**Supplementary Fig. S9**). To confirm the SBS.A3G signature we identified, we employed two additional orthogonal analytical methods for mutational signature extraction that rely on different mathematical procedures. The SigneR package(44) provides a full Bayesian treatment to the non-negative matrix factorization (NMF) model. The HDP package(45) utilizes the hierarchical Dirichlet process (**Methods**). Both methods identified a mutational signature that was present in the *hA3G*(+) *mA3*(−/−) tumors but absent from *mA3*(−/−) tumors consistent with the SBS.A3G signature (Cosine similarity >0.7). The extracted signatures from the three methods confirmed the key defining features of SBS.A3G, including the predominance of C>T mutations in CCC, CCT, and TCC motifs (**Supplementary Fig. S10A and S10B**).

We then compared SBS.A3G to a previously described mutational signature induced by transgenic expression of human APOBEC3A in mice (SBS.A3A)(11) (**Methods**). SBS.A3G showed low cosine similarity to SBS.A3A, which is characterized by predominant C>T substitutions in the TCA and TCT motifs (**Fig. 3D and 3E**). Furthermore, SBS.A3G showed a low cosine similarity to the mutational signature detected in C57BL/6J mice expressing wild-type mouse *Apobec3* (SBS.mA3) (**Supplementary Fig. S8C-E**). Finally, we characterized the relationship between the SBS.A3G signature and the Catalogue Of Somatic Mutations in Cancer (COSMIC) single base substitution signatures derived from human cancers (**Methods**). We confirmed that SBS.A3G had a low cosine similarity to SBS2 and SBS.A3A, which are characterized by C>T substitutions predominantly in the TCW motifs (**Fig. 3F; Supplementary Fig. S10C**). Together, these data suggest that APOBEC3G induces a distinct mutational signature from other APOBEC3 family members.

### APOBEC3G contributes to mutagenesis in human cancers

We asked whether the SBS.A3G signature we identified contributes to the mutational profiles of human cancers. We reasoned that APOBEC3G expression is a prerequisite for APOBEC3G-induced mutagenesis. We examined the mRNA expression of *APOBEC3G* and the resulting mutational signatures in 8292 tumors from 21 cancer types from The Cancer Genome Atlas (TCGA) pan-cancer cohorts(46,47). *APOBEC3G* mRNA was ubiquitously expressed in all tumor types (mean RNA-seq by Expectation-Maximization (RSEM) value: 368.7, mean expression ratio normalized to the mRNA expression of the housekeeping gene TATA-Box binding protein (*TBP*): 1.46) (**Fig. 4A**). Diffuse large B-cell lymphoma (DLBC), urothelial bladder carcinoma (BLCA), and renal clear cell carcinoma (KIRC) had the top normalized *APOBEC3G* to *TBP* expression ratios (4.9, 3.3, 3.1, respectively). Furthermore, the median C>T substitution counts in the CCC, CCT, and TCC, constituting the SBS.A3G component motifs, significantly correlated with the median *APOBEC3G* mRNA expression ratio (R=0.49, *P*=0.02) (**Fig. 4B; Supplementary Fig. S12A**). To exclude the possibility that APOBEC3 expression in bulk tumor RNA sequencing originated from the infiltrating immune cells, we examined the expression level of APOBEC3G in cancer cell lines in the Cancer Cell Line Encyclopedia (CCLE) data(48,49). We identified significant *APOBEC3G* mRNA expression in the cancer cell lines (**Supplementary Fig. S11**).

**Figure 4.**
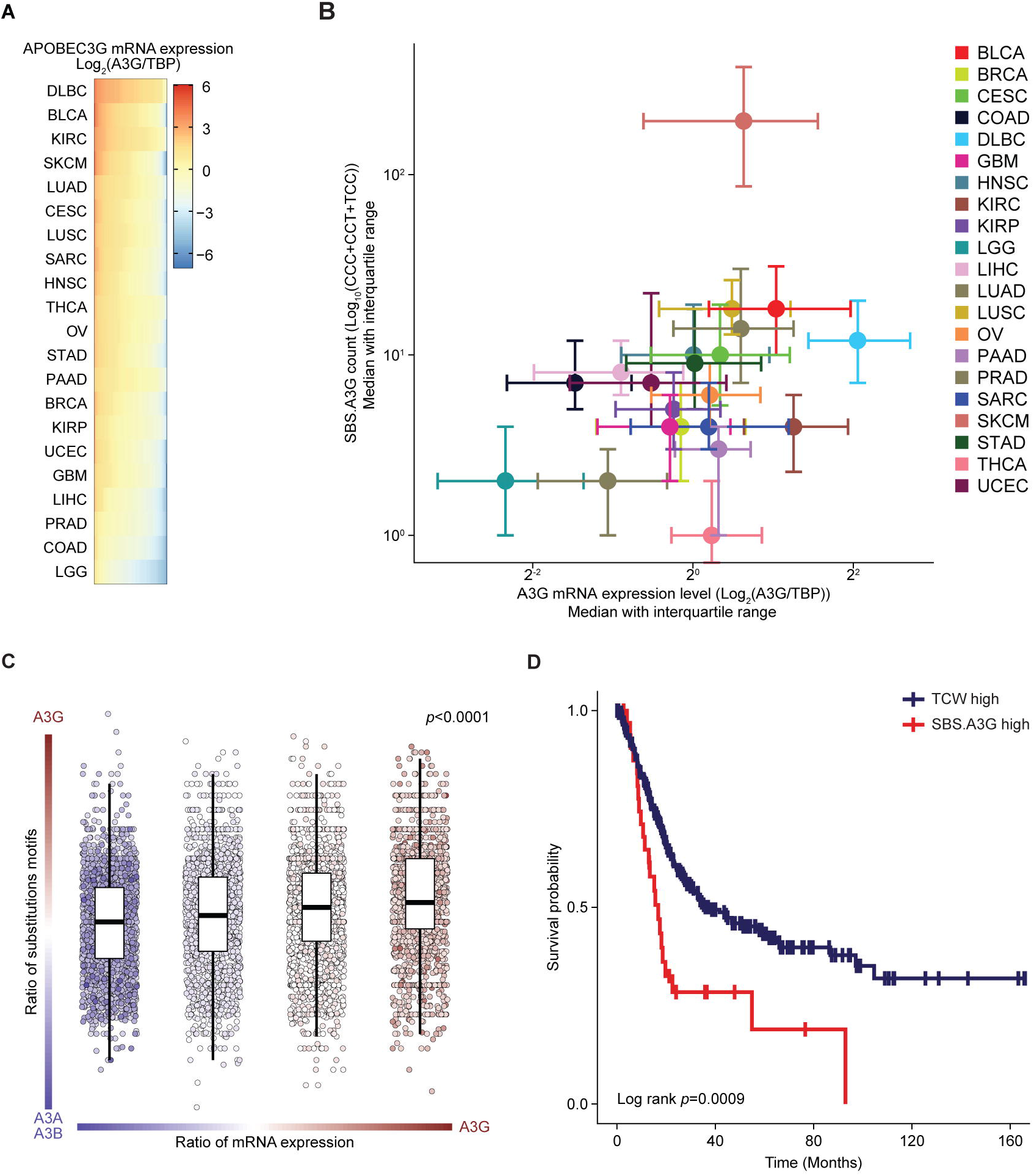
The mutational impact of APOBEC3G in human cancers. A) mRNA expression level of *APOBEC3G* (normalized to *TBP*) in different cancer types in the TCGA pan-cancer cohort. The scale bar indicates the *APOBEC3G* mRNA level normalized to *TBP*. B) Correlation between *APOBEC3G* mRNA expression and SBS.A3G mutational loads in TCGA pan-cancer cohorts. Nonparametric Spearman correlation analysis was performed between the median *APOBEC3G* mRNA expression and median C>T substitution counts in CCC, CCT, and TCC motifs. R=0.49, *P*=0.02. Each dot represents the median *APOBEC3G* mRNA expression level (normalized to *TBP*) (horizontal) and the median C>T counts in CCT and TCC motifs (vertical) with error bars representing IQR. C) *APOBEC3G* mRNA expression positively correlates with the percentage of C>T substitutions in the CCC, CCT, and TCC motifs attributed to APOBEC3G. It negatively correlates with the percentage of C>T substitutions in TCW motifs attributed to APOBEC3A and APOBEC3B in TCGA pan-cancer cohorts. The *P* value was calculated by Kruskal-Wallis nonparametric test. The box plot shows the ratios as median with IQR without outliers. The lower whisker indicates Q1-1.5*IQR. The upper whisker indicates Q3+1.5*IQR. Each dot indicated one sample. The dot color presented the ratio of mRNA. D) Patients in the SBS.A3G-predominant group (CCC, CCT, and TCC) had lower survival than patients in the TCW-predominant groups, irrespective of the total mutational burden in the TCGA bladder cancer cohort. Log-rank test. TBP: TATA-Box binding Protein. IQR: interquartile range. A3A: APOBEC3A. A3B: APOBEC3B. A3G: APOBEC3G. TMB: Tumor mutational burden. Abbreviations for the TCGA cancer types are available at https://gdc.cancer.gov/resources-tcga-users/tcga-code-tables/tcga-study-abbreviations.

We then used three different fitting pipelines, DeconstructSigs (DS)(50), sigLASSO (SL)(51), and MutationalPatterns (MP)(52), to estimate the contribution of mutation counts attributed to each COSMIC signature in each tumor from TCGA pan-cancer with all COSMIC exon signatures (**Methods**). We found that in the urothelial bladder carcinoma (BLCA) cohort, 13% of patients harbor SBS.A3G with a median of 17 contributing counts (**Supplementary Fig.S13A)**. The median counts and proportion of tumors with contributions from different TCGA cohorts were averaged based on the highly consistent results from these three different fitting approaches (**Supplementary Fig. S13A and S13B)**.

We then set out to dissect the relative contributions of the APOBEC3A, APOBEC3B, and APOBEC3G proteins, which are frequently co-expressed in a given tumor(53–55). We examined the correlation between the ratio of *APOBEC3G* to *APOBEC3A* and *APOBEC3B* mRNA expression and the fraction of C>T substitutions in the motifs preferred by each protein. We discovered a significant correlation between the *APOBEC3G*/(*3A*+*3B*) mRNA ratio and the C>T substitution ratio in (CCC+CCT+TCC)/(TCA+TCT) (*P*<0.0001) (**Fig. 4C**). These data indicate that APOBEC3G-induced mutagenesis is independent of APOBEC3A and APOBEC3B. To understand the impact of APOBEC3G-induced mutations on clinical outcomes, we assigned urothelial bladder cancer patients in the TCGA cohort into SBS.A3G-predominant (CCC+CCT+TCC) and TCW-predominant subgroups based on the preferred motifs by each protein. Patients in the SBS.A3G-predominant subgroup had lower survival than patients in the TCW-predominant subgroup regardless of the total mutational burden (log-rank *P*=0.0009) (**Fig. 4D; Supplementary Fig. S14A**). These data confirm that the negative impact of APOBEC3G on survival we initially observed in our bladder cancer mouse model extends to bladder cancer patients.

## Discussion

Mutational signatures in human cancers have been associated with multiple mutagenic processes(56–60). However, human cancers are intrinsically noisy systems harboring a continuous interplay of mutagenic and DNA repair processes(14). Consequently, approaches relying on the analysis of sequencing data from human cancers, followed by heuristic attribution of mutational signatures to a particular mutagenic process, are inherently limited(12–14). These challenges have hindered our understanding of the specific contributions to mutagenesis of each of the seven APOBEC3 proteins, which are frequently co-expressed in human cancer cells(53–55). Here, we focused on APOBEC3G, a cytidine deaminase that restricts lentiviruses by mutating the viral genome(24,36,38,61). However, whether it can mutate genomic DNA in cancer cells has remained unknown until our study. To dissect the mutagenic impact of a single member of the human *APOBEC3* family in an experimentally controlled bladder cancer mouse model, we leveraged the differences between mouse and human *APOBEC3* loci, which are separated by 76 million years of evolution(62).

In our examination of the TCGA pan-cancer dataset, we observed significant levels of *APOBEC3G* expression across several cancer types. However, determining a definitive causal relationship between the mRNA expression of individual *APOBEC3* genes and their respective signatures in human cancer genomes is challenging for several reasons.

First, mutational signatures can be imprinted on the nuclear genome cumulatively over long-time intervals or in punctuated episodes. Therefore, the expression level of a given mutagenic protein at the time of sampling may not reflect its expression level at the time of mutagenesis in human cancers(63–65). Second, different APOBEC3 proteins may be expressed in the same cancer cell simultaneously, making it challenging to attribute specific mutational signatures to individual APOBEC3 proteins. Here, using a model with transgenic expression of human APOBEC3G on a null mouse *Apobec3* background, we provide cause-and-effect experimental evidence showing that human APOBEC3G directly contributes to mutating cancer genomes *in vivo*. Our data show that APOBEC3G induces genomic instability, increasing tumor mutational burden, copy-number loss events, and clonal diversity. Clonal diversity is thought to arise from the acquisition of somatic mutations and chromosomal structural variants(30,66), and high intra-tumoral heterogeneity was previously associated with poor cancer outcomes and the evolution of drug resistance(32,67–69). We also discovered that APOBEC3G significantly increased kataegis in a pattern consistent with the known processive nature of its catalytic deamination effects(5,6,25–27). Kataegic events are enriched in genomic rearrangements(70,71) and chromothripsis regions(26,72) in several cancer types. These processes contribute to oncogene amplification or the inactivation of tumor-suppressing genes, which are important driver events in human cancers(72). Longitudinal computational analyses of the mutational profiles of serial tumors from individual patients show that mutational signatures attributed to APOBEC3 proteins (SBS2 and 13) are associated with subclone expansions in several cancer types, including urothelial bladder cancer(4,50,73–77). Ongoing efforts to develop small molecule inhibitors of APOBEC3 proteins to restrict cancer evolution require specific knowledge of the role of the individual APOBEC3 proteins responsible for driving intra-tumoral heterogeneity(3,24).

Although APOBEC3G-induced hypermutations in the HIV genome have been described(18,36), the signature attributed to APOBEC3G in the cancer genome has not been previously revealed by pan-cancer studies(1,5), even for cancers in patients with HIV infections. However, APOBEC3G expression was shown to be both upregulated(78,79) and downregulated(80,81) following HIV infection. Immune cells, rather than epithelial cancer cells, are the preferred targets for HIV *in vivo*. Thus, non-hematological malignancies in HIV-positive patients are not expected to have a heavy burden of APOBEC3G-induced mutations. Here, we adopted an experimental approach to establish the APOBEC3G-induced signature using three well-established orthogonal computational frameworks(12,13,44,45). The signature we identified (SBS.A3G) includes C>T substitutions in the CC motif previously reported in the viral genome(18,36) and substitutions in the TCC motif consistent with the previously identified secondary APOBEC3G’s preference for the TC motif in *in vitro* assay(82) and the viral genome in cell lines(18,36,42,83). Our findings suggest that the motifs preferred by APOBEC3G in the HIV and nuclear genomes overlap.

The subcellular distribution of APOBEC3G is predominantly cytoplasmic(84). However, our data suggest that APOBEC3G could access the nuclear compartment of bladder cancer cells, which is consistent with previous studies that found small amounts of APOBEC3G are present in the nuclei of human T lymphocyte lines(85,86) and that APOBEC3G significantly contributes to an increase in DNA damage and genomic instability in multiple myeloma cell lines(87). It is important to note that the mechanisms resulting in the mutations induced by APOBEC3G in cancer genomes are not entirely dependent on the motif binding preferences of the enzyme. Factors exclusive to the nuclear genome may alter the likelihood of cytidine deamination, including interactions between APOBEC3G and single-strand DNA binding proteins(88,89) and epigenetic modifications of cytidine, which render them more resistant to APOBEC3G-induced deamination(90–92). These factors collectively cooperate to determine the final signature imprinted by APOBEC3G on genomic DNA.

SBS.A3G had low cosine similarity to motifs attributed to other proteins in the APOBEC3 family. The differences in motif preferences and mutational patterns between APOBEC3G and APOBEC3A and APOBEC3B potentially explain the differences between our experimentally induced SBS.A3G and the COSMIC SBS2 and SBS13 signatures, which are classically associated with APOBEC3A and APOBEC3B(1,2,5,71). In addition, we found that bladder patients in the group dominated by APOBEC3G-induced mutational signatures had shorter overall survival than those enriched in signatures attributed to APOBEC3A and APOBEC3B, consistent with a previous report(93). These data suggest that individual APOBEC3 proteins impact clinical outcomes differently. It is important to note that in non-muscle invasive bladder cancers, non-small cell lung, and breast cancers, APOBEC3 mutations are collectively associated with poor clinical outcomes(94–96). Our study has some potential limitations, one of which is the need for an initial carcinogenic event in our mouse model. We chose to use BBN, an alkylating nitrosamine, similar to cigarette smoke carcinogens which are implicated in human urothelial bladder cancer(97). This model recapitulates human urothelial bladder cancer but notably lacks APOBEC3-induced mutational signatures(97).

In summary, we show that APOBEC3G contributes to mutagenesis, kataegis, and intra-tumor heterogeneity in cancer genomes. Our findings potentially inform future therapeutic efforts to restrict tumor evolution by targeting specific APOBEC3 enzymes(3).

## Methods

### Animals

*Apobec3* null mice (*mA3*(−/−)) and mice expressing the human *APOBEC3G* transgene on a null *mApobec3* (*hA3G*(+) *mA3*(−/−)) were previously described(21,22). A breeding scheme was used in which one parent carried the transgene (*hA3G*(+) *mA3*(−/−)), and the other parent did not (*mA3*(−/−)). This allowed us to use the *mA3*(−/−) littermates as the control group in our study. Mice were weaned at three weeks of age and group-housed. Genotyping of each animal was performed by Transnetyx using real-time PCR to detect the human transgenes and confirm the absence of *mApobec3* using primers as previously described(21,22). The wild-type C57BL/6 male mice as control were purchased from Jackson Laboratory. The animal experiments were carried out following Weill Cornell Medical College Institutional Animal Care and Use Committee guidelines (IACUC Protocol 2017-0048).

### Cell lines

The HEK293T and 5637 cell lines were purchased from ATCC (CRL-3216™, HTB-9™). The RT112/84 cell line was purchased from Millipore Sigma (#85061106). The HEK293T cells were incubated in a humidified constant 37C and 5% CO2 incubator in DMEM medium (Gibco) with 100U/ml Penicillin-Streptomycin (Gibco) and 10% FBS (Omega Scientific). The 5637 and RT112/84 were incubated in a humidified constant 37C and 5% CO2 incubator in RPMI-1640 (Gibco) and EMEM (ATCC) medium with 100U/ml Penicillin-Streptomycin (Gibco) and 10% tetracycline-free FBS (Omega). The cells were tested for *Mycoplasma* contamination with the PCR detection kit (ABM) according to the manufacturer’s instructions.

### Study design

N-butyl-N-(4-hydroxybutyl)nitrosamine (BBN) was purchased as a single batch from TCI America (Portland, Oregon, USA, Batch ODW3F-FH). Red tinted water bottles (ANCARE) were used during drug administration since BBN is a light-sensitive compound. *Ad libitum* BBN was administered at a concentration of 0.05% in water and replenished twice weekly. Bottles were weighed to determine the volume drunk pre-cage during each period. The volume consumed by each animal was determined as the quotient of total water drunk per cage to the number of animals in that cage. Mice began BBN administration via drinking water between 8-10 weeks and continued for 12 weeks. After 12 weeks of *ad libitum* carcinogen administration, mice were returned to unaltered drinking water for 18 weeks to provide adequate time for APOBEC3G-induced mutagenesis. All experimenters were blinded to the *mA3*(−/−) and *hA3G*(+) *mA3*(−/−) genotype due to the mixed genotype populations in each cage. The wild-type C57BL/6J mice were treated in separate cages. Survival curves were calculated as the percentage of mice that expired prior to the 30-week time point.

### RNA extraction and qPCR analysis

The bladder, lung, and spleen tissues were harvested from three *hA3G*(+) *mA3*(−/−) and *mA3*(−/−) mice, respectively. Tissues were snap-frozen using liquid nitrogen. Tissues were homogenized using BioMasher II (TaKaRa) with RLT Plus buffer (1% β-mercaptoethanol).

Total RNA was extracted using the RNeasy Plus Mini Kit protocol (Qiagen). RNA concentration was determined by NanoDrop (ThermoFisher Scientific). cDNA was synthesized using the SuperScript III First-Strand Synthesis System (Invitrogen). Real-time PCR was performed by LightCycler 480 (Roche) using Power SYBR Green Master Mix (Applied Biosystems). All reactions were run in triplicate and analyzed using the LightCycler 480 software. *APOBEC3G* expression data were normalized to *GAPDH*. The qPCR primers were listed in **Supplementary Table S4**.

### SDS-PAGE Western blotting

Whole bladder protein lysate was obtained by homogenizing bladder tissue using Biomasher II (TaKaRa) in ice-cold RIPA buffer with the protease inhibitor (ThermoFisher Scientific). Protein concentration was determined using a Pierce BCA Assay (ThermoFisher Scientific). The lysate was run on a 10% SDS page gel with MOPS buffer and transferred using the iBlot system (ThermoFisher Scientific). The anti-Myc antibody (Abcam, ab9106, 1:1000) and anti-α-tubulin (Millipore, 05-829, 1:1000) were separately used as primary antibodies at 4C overnight. HRP conjugated secondary antibodies (Goat anti-Rabbit, 32260, and Goat anti-Mouse, 32230, Invitrogen, 1:1000) were incubated for 1hr at room temperature. Blots were developed using Luminata Forte Poly HRP Substrate (Millipore) reagent and imaged on the ChemiDoc imager (BioRad). The protein lysate from the wild-type HEK293T cell and HEK293T transfected with the hAPOBEC3G-Myc-T2A-eGFP vector using Lipofectamine2000 (ThermoFisher Scientific) were used as negative and positive controls. This vector was synthesized by VectorBuilder, and its sequence was validated by Sanger sequencing.

### Generation of doxycycline-inducible cell lines

The doxycycline-inducible GFP-tagged APOBEC3G vector and PiggyBac transposase vector were obtained as a gift from Dr. John Maciejowski Lab. The sequences were validated by Sanger sequencing. The human urothelial bladder tumor cells were transfected with the doxycycline-inducible GFP-tagged APOBEC3G vector and the PiggyBac transposase vectors with the Lipofectamine2000 transfection agent (ThermoFisher Scientific). G418 was used for the selection of the clones with stable genomic insertion of the GFP-tagged APOBEC3G component. The serial dilution method isolated single-cell clones in 96 well plates. The expression of APOBEC3G in each single-cell clone was validated by SDS-PAGE western blot.

### Nuclear and cytoplasmic extraction

The NE-PER™ nuclear and cytoplasmic extraction kit (ThermoFisher Scientific) was used according to the manufacturer’s instructions to isolate nuclear and cytoplasmic protein fractions from the cells after two days of doxycycline induction. The anti-GFP antibody (Abcam, ab13970,1:2000), anti-α-tubulin (Millipore, 05-829,1:1000), and anti-Lamin A/C (Cell Signaling Technology, 4777S,1:1000) were used as primary antibodies for 2hr at room temperature. The anti-Myc antibody (Abcam, ab9106,1:1000) was used as the primary antibody at 4C overnight. HRP conjugated secondary antibodies (Goat anti-Rabbit, 32260, Rabbit anti-Chicken, 31401, and Goat anti-Mouse, 32230, Invitrogen, 1:1000) were used for 1hr at room temperature. Blots were developed using Luminata Forte Poly HRP Substrate (Millipore) reagent and imaged on the ChemiDoc imager (BioRad).

### Immunofluorescence (IF) microscopy

The cells were plated in a T25 flask and treated with vehicle (DMSO) or doxycycline for 48hrs. Cells were harvested, and nuclei were isolated using hypotonic buffer as previously described(98). The isolated nuclei were spun down onto the coverslip precultured with poly-L-ornithine (Sigma) at 400G for 5 mins, then fixed by 4% paraformaldehyde at room temperature for 10 mins, following 3 times washing with PBS. After slides were made, the IF protocol was performed. The fixed nuclei were treated with 0.5% Triton X-100 in PBS, then blocked by 2.5% normal goat serum in PBS. The anti-α-tubulin antibody (Millipore, 05-829,1:1000) was used as the primary antibody at 4C overnight. Then, the AlexaFluor-594 goat anti-mouse antibody (Invitrogen, a11032, 1:1000) was used as a secondary antibody for 2hr at room temperature. After that, the slide was sealed with ProLong Gold Antifade Mountant with DAPI (Invitrogen, p36931). High-resolution imaging was performed using a DeltaVision Elite system, with a 60X/1.42 oil objective (Olympus) and EDGE sCMOs 5.5 camera. 12 different fields of view for each group were randomly picked for imaging. Different channels were acquired for DAPI, FITC, and TRITC filters. Z-stacks of 0.2um increments for a total of 20 stacks (4um) were acquired per field of view. The captured images for the nuclei were adjusted and quantified using ImageJ Fiji software as follows. The Z stacks were summed to create a sum projection, and background subtraction was performed. Then the image was split into three channels, DAPI (blue), GFP (green), and α-Tubulin (red). The DAPI image was processed by Gaussian blur, auto-threshold using the Otsu method, and then converted into a mask to create a list of regions of interest based on size and circularity. The manual correction was performed to ensure that only single nuclei were quantified. Next, the mean intensity value of the GFP (green) channel for regions of interest was measured. Multiple comparisons were performed for statistical analysis.

### Confocal laser scanning fluorescent microscopy and 3D image reconstruction

A single isolated nucleus fluorescent image was captured using Zeiss LSM880 inverted confocal microscope with a 63X/1.4 oil Dix M27 objective. The DAPI and FITC channels were scanned using an auto-optimized setup by microscopy software, ZEN Black 2.1 SP3 LSM. The image data was imported into the Imaris software (Oxford instruments) for 3D reconstruction. The nuclear surface was reconstructed based on the DAPI staining. Then the GFP signal within the nuclear surface was isolated and reconstructed by default. The green color represents the GFP signal, and the violet represents the reconstructed nuclear surface.

### Excision and processing of bladder tumors

Mice were euthanized at the 30-week pre-specified timepoint. The bladders were excised, weighed, and measured using a pathology ruler. Bladders were bisected along the sagittal plane, placed urothelium-side down in a cryomold, embedded in OTC (Tissue-Tek) using 2-methylbutane with dry ice, and sectioned with a cryostat. Bladders from expired mice were extracted, and cryomolds were created. Cryomolds were stored at −80C. Frozen slides were generated and converted to FFPE slides for H&E staining at the Department of Pathology at Weill Cornell Medicine for histopathological review. The H&E slides underwent histopathological review by a fellowship-trained genitourinary pathologist (FK) to annotate cancer-involved regions and determine the tumors’ extent. The pathologist was blinded to the genotype and experimental condition of the samples during the review.

### Generation of murine tumor organoids

After excision of the bladder, a piece of tissue was obtained from the bladder tumor and placed in Advanced DMEM (Gibco) containing collagenase IV (100U/ml) (Gibco) and 10uM ROCK inhibitor Y-27632 (Selleckchem, S6390) at 37C for 6 hrs. After the cell solution was filtered by cell stainer (FALCON), the tumor cells were harvested, resuspended in an organoid media with Matrigel (Corning), and plated as 3D droplets in a 6-well suspension plate (Cellstar). The organoid media composition was previously described(99). After sufficient growth in the 3D culture, the cells were passaged into the 2D culture in the same media.

### DNA extraction, library preparation, and whole-exome sequencing

Frozen tumor tissue was punched on dry ice using a 1mm biopsy punch (Miltex) based on the region marked by the pathologist. The DNA from mouse bladder tumors and matched germline DNA from mouse tails were extracted using the DNeasy Blood Tissue kit (Qiagen) according to the manufacturer’s instructions. DNA was stored at −80C until library preparation. Quality control for DNA samples was performed using TapeStation (Agilent), and libraries for Whole Exome Sequencing were prepared with the Agilent SureSelect kit (SureSelect Mouse All Exon Kit) at the Weill Cornell Medical College genomics core facility. Pooled samples were loaded on NovaSeq6000 (Illumina) and sequenced (paired-end 2X100). The sequencing coverage of bladder tumors and matched germline tissues were generated using Picard before removing PCR-duplicate reads.

### Alignment and somatic variant calling

Mouse reference genome GRCm38/mm10 was used for short reads alignment using our in-house alignment pipeline. Short reads were trimmed for adapter sequences, aligned by BWA MEM, indel realignment was performed via GATK, and deduplication of PCR duplicates was performed by Picard Tools. Somatic single nucleotide variants (SNVs) were called using TCGA MC3 Variant Calling Strategy, which merges the seven proven-performed variant calling methods, including Strelka2, MuSE, MuTect2, Pindel, RADIA, SomaticSniper, and VarScan, where MuTect2 and Strelka2 replaced MuTect and Strelka(100). Default parameters were used for these seven tools, except for instances identified by the original MC3, where non-default settings yielded optimal performance. MuSE was implemented with the germline resource. Pindel was implemented with a blacklist reference made available through the ENCODE Blacklist resource. Post-hoc filtering of the variants identified by each caller was done identically to the MC3 pipeline except as follows. For MuSE, variant calls were retained at least Tier 4-level significance, which corresponds to the calls with an added false positive and false negative probability of less than 1%. For Strelka2, the variant calls were filtered to those that passed the tool’s internal significance testing. For Mutect2, the variant calls were filtered in two steps, with the first entailing filtering based on the significance scoring of each call and the estimation of sample contamination. These methods were implemented via the *CalculateContamination* and *FilterMutectCalls* method in GATK4. The second step consisted of quantifying nucleotide substitution errors caused by mismatched base pairings during various sample/library preparation stages. Such errors include artifacts introduced before the addition of adapters and those introduced after target selection. These sequencing artifacts were then filtered from the original Mutect2 calls using the *CollectSequencingArtifactMetrics* and *FilterByOrientationBias* methods of GATK4. All filtered variant calls from each tool were sorted using *vcf-sort* from the vcftools package and standardized as in the MC3 pipeline. After standardization, the calls from multiple tools were merged into a single file. The merged variant calls were further filtered for sequence artifacts described above and calls from regions in the capture region designed by SureSelect. Post-filters included the variants from multiple callers 1) At least two callers should call the variants. 2) Only the single nucleotide variants (SNVs) were included for further analysis. 3) The variants were filtered by mouse dbSNP (built 146 for mouse mm10 assembly). 4) The total read counts for germline ≥ 20 and tumor ≥ 40. 5) Alt read counts in germline < 5 and the ratio of tumor variant allele frequency (VAF)/germline VAF ≥ 5. 6) The alt read counts in tumor > 10. Total mutations per Mb were calculated as the total number of somatic variants, including the missense and nonsense in the coding region after annotation, divided by 40.6 Mb pair, the protein-coding region size.

### Annotation of somatic calls

Somatic calls from the pipeline above were annotated using VEP version 95 and converted to MAF (Mutation Annotation Format) format using the vcf2maf.pl script (https://github.com/mskcc/vcf2maf).

### Copy number variant analysis

The copy number variants (CNV) were evaluated using CNVkit(29). The CNV results were visualized as the heatmap of binned log2 ratios using the R/ComplexHeatmap package(101) and circus plots using the R/RCircos package(102). The cytoband information and known gene definitions of mm10 assembly were extracted using R/UCSC.Mouse.GRCm38.CytoBandIdeogram,R/TxDb.Musculus.UCSC.mm10.knownGene and R/org.Mm.eg.db packages. The log2 value < −0.2 was considered copy number loss, and log2 value > 0.2 was considered copy number gain. The length of segments involved with copy number variants was then calculated.

### Clonal diversity analysis

Clone numbers and distribution were analyzed using the Pyclone-vi computational tool(33) and EXPANDS tool(31) using SNVs and CNVs data as inputs. The CNV data inputs for Pyclone-vi were generated by CNVkit. The CNV data inputs for EXPANDS were generated by the TITANCNA package(103). True diversities (Hill numbers, H) were calculated as 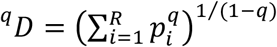. Shannon entropy (*H′*) was calculated as *In*(^1^*D*). Diversity profiles for the *hA3G*(+) *mA3*(−/−) and *mA3*(−/−) tumors were generated by graphing H(p,q) against q, where q is the index’s sensitivity parameter q. Here, ^1^*D* is the Hill number of order 1. Differences between the medians of Hill numbers between the *hA3G*(+) *mA3*(−/−) and *mA3*(−/−) tumors were separately analyzed using bootstrap-coupled estimation statistics and visualized as Gardner-Altman plots(104). The bias-corrected and accelerated 90% CI for the differences in medians were calculated from 5000 bootstrapped samples. The *P* value reported is the likelihood(s) of observing the effect size(s) if the null hypothesis of zero difference is true. Phylogenetic trees were generated by EXPANDS and visualized using the ggtree package(105,106).

### Kataegis analysis

Clustered mutational events were identified using two approaches. First, we defined kataegic clusters by an inter-mutational distance (<1Kb) and mutation number (≥ 4) within a cluster. The distance and the genomic location of somatic mutations were generated using MutationalPatterns. The second method, SeqKat, identifies the clustered mutational events based on the binomial test(28). APOBEC-related kataegic events were defined as mutational clusters harboring enrichment of C>T mutations *in cis* based on the binormal test (at least 2 C>T mutations *in cis* in 4-5 clustered mutations or at least 3 C>T mutations *in cis* in 6-10 clustered mutations). Fisher’s exact test was used to compare the number of APOBEC-related kataegis within the total kataegic clusters in the *hA3G*(+) *mA3*(−/−) and *mA3*(−/−) tumors and compare the enrichment of C>T substitutions *in cis* compared to other C>T substitutions (singlet or clustering *in trans*) within kataegic loci between the *hA3G*(+) *mA3*(−/−) and *mA3*(−/−) tumors.

### Transcriptional and replicative strand asymmetry of APOBEC3-induced mutations

The transcriptional strand annotation for each mutation was generated using the mut_strand function in MutationalPatterns. The ratio of the total count of C>T mutations in the transcribed strand/the total count of C>T mutations in the un-transcribed strand were calculated for each tumor. The ratios of *hA3G*(+) *mA3*(−/−) tumors were standardized to the mean ratio of the *mA3*(−/−) tumors. For the replicative strand bias analysis, the reference strand annotations were obtained from Riva *et al*.(45). Then, the replicative strand information for each mutation was generated using the mut_matrix_stranded replication model in MutationalPatterns. The replicative strand ratios of C>T mutations were then calculated for *hA3G*(+) *mA3*(−/−) tumors, then standardized to *mA3*(−/−) tumors.

### Principal component analysis

Trinucleotide mutational context for each sample was generated by MutationalPatterns(52). Based on the trinucleotide context for each sample, the PCA analysis was performed by factoextra package. The confidence ellipse for each group represents the 95% confidence interval. Permutational Multivariate Analysis of Variance test was used to calculate the *P* value via vegan package.

### Mutational signature analysis

We employed a bootstrap resampling method(12,13) to extract the net APOBEC3G-induced mutational signature. Apart from the transgenic expression of human APOBEC3G, the backgrounds of the two strains of mice are genetically identical. Compared with the *mA3*(−/−) mice, we hypothesized that the *hA3G*(+) *mA3*(−/−) tumors harbor human APOBEC3G-induced mutational signatures, which increased the total mutational burden. To detect the mutational shift distance (MSD) between the *hA3G*(+)*mA3*(−/−) and *mA3*(−/−) tumors, we used bootstrap resampling as previously described(12). The normalized distance was calculated between the centroids of resamples and the original samples. The centroids were determined based on the mean value of the percentage of substitutions in the 96 channel matrices. The distribution of the normalized distance was obtained by repeating the previous step 10,000 times. The threshold distance for each group corresponding to a *P* value (*P*=0.05) was calculated. A significant mutational spectrum shift between the *hA3G*(+) *mA3*(−/−) and *mA3*(−/−) tumors was determined to be present if it crossed the threshold of the distances of two resampling groups(13). To detect significantly increased substitutions (*P* value <0.1) and extract the distinct APOBEC3G signature, the centroid of bootstrapped the *hA3G*(+) *mA3*(−/−) tumors was compared to the centroid of the *mA3*(−/−) tumors. For significantly increased substitution types, the magnitude of the increased counts was calculated as the difference between the centroids multiplied by the mean counts of resampled tumors. This step was then repeated 10,000 times. The significantly increased counts in each trinucleotide context were averaged to construct the APOBEC3G signature *de novo* as previously described(12). To validate the signature we extracted, we use two additional methods for signature extraction using the SigneR(44) tool, which provides a full Bayesian treatment to the non-negative matrix factorization (NMF) model, and the HDP package(45), which uses the hierarchical Dirichlet process to *de novo* extract mutational signatures. For each approach, the trinucleotide mutational matrices of *mA3*(−/−) and *hA3G*(+) *mA3*(−/−) tumors were separately imported to *de novo* extract the signatures. After extraction, the cosine similarity was calculated between the signatures from *mA3*(−/−) and *hA3G*(+) *mA3*(−/−) tumors by MutationalPatterns. Then, the unique signature from *hA3G*(+) *mA3*(−/−) tumors, which had low cosine similarity to the signatures extracted from the *mA3*(−/−) tumors, was considered the signature induced by APOBEC3G (SBS.A3G).

### Mutational signature induced by transgenic APOBEC3A expression

The raw sequencing data from 5 tumors from transgenic APC^min^ C57BL/6J mice expressing human APOBEC3A and 4 tumors from transgenic APC^min^ C57BL/6J mice were downloaded from the SRA database (BioProject ID: PRJNA655491)(11). We reanalyzed the data and performed mutational calling using our MC3 pipeline with the post-filters as below: 1) At least two callers should call the variants. 2) Only the single nucleotide variants (SNVs) were included for further analysis. 3) The total tumor read counts ≥ 20. 4) The normal VAF ≤ 0.01. 5) The tumor VAF > 0.02. The trinucleotide mutational context for each sample was generated by MutationalPatterns.

### Cosine similarity to COSMIC signatures

The cosine similarity between the experimentally-extracted signatures and Catalogue Of Somatic Mutations in Cancer (COSMIC) signatures (sigProfiler_exome_SBS_signatures) was computed using MutationalPatterns. The signatures extracted from whole-exome sequencing of murine tumors were normalized for the trinucleotide frequency in the human exome. The trinucleotide count in the mouse SureSelect whole-exome sequencing region was calculated using the triCount.R function in the mutationalProfiles package. The trinucleotide count of the human GRCh37/hg19 whole exome was obtained from the DeconstructSig package. The adjusted SBS.A3G was then used to calculate the cosine similarity to COSMIC exome signatures V3.

### Fitting extracted signatures to human cancer

Three different pipelines were used to fit the experimental signatures to human cancer. 1) MutationalPatterns, which is a backward method to find a non-negative linear combination of mutation signatures to reconstruct the mutation matrix (fit_to_signatures_strict function). 2) deconstructSigs, which uses a multiple linear regression model to reconstruct the mutational profile of a single tumor sample(50). 3) sigLASSO, which jointly optimizes the likelihood of sampling and signature fitting(51). Given the overlap between the UV signatures (SBS7a and SBS7b) with the experimentally validated SBS.A3G, we substituted SBS7a and SBS7b with SBS.A3G to the 63 other COSMIC V3 exon signatures to fit the signature in patients’ tumors in UV-unrelated tumors. The consistency of the median proportion of patients harboring contribution from the respective signature and the signature’s contributing counts among three fitting pipelines were measured by Pearson correlation analysis using the corrplot R package.

### Analysis of APOBEC3-preferred motifs in the HIV genome

The data was obtained from Dr. Linda Chelico(43). APOBEC3G-induced substitutions in the *pol* gene of HIV were measured in an isogenic HEK293T cells system expressing APOBEC3G(43). We reanalyzed the data and calculated substitutions based on the probability of the substitution at each position within the sequence context. First, we set up a filter for variant allele frequency of 0.01 (relevant alteration read count >100) to exclude background noise and call high-confidence substitutions at each position. Then, the counts for C>T substitutions in TC and CC motifs were calculated.

### Statistical tests

One-tailed Mann-Whitney U test for continuous variables, two-tailed 2X3 table Chi-Square test or Fisher’s exact test for categorical variables, Kruskal-Wallis nonparametric test for multiple groups, nonparametric Spearman correlation, and log-rank test for survival analysis was performed using GraphPad Prism version 8.4.3 statistical analysis software. *P*<0.05 was considered statistically significant.

## Ethical Approval

All animal experiments were carried out following Weill Cornell Medical College Institutional Animal Care and Use Committee guidelines (IACUC Protocol 2017-0048).

## Data availability statement

The TCGA pan-cancer human cancer data are available at cbioportal and dbGaP under the accession number PHS000178.

## Code availability statement

The open source code used in this paper were listed : BWA MEM (https://github.com/lh3/bwa), GATK (https://github.com/broadinstitute/gatk), Picard (https://github.com/broadinstitute/picard), Strelka2 (https://github.com/Illumina/strelka), MuSE (https://github.com/danielfan/MuSE), MuTect2 (https://github.com/broadinstitute/gatk), Pindel (https://github.com/genome/pindel), RADIA (https://github.com/aradenbaugh/radia), SomaticSniper (https://github.com/genome/somatic-sniper), VarScan (https://github.com/dkoboldt/varscan), VEP (https://github.com/Ensembl/ensembl-vep), vcf2maf (https://github.com/mskcc/vcf2maf), SeqKat (https://github.com/cran/SeqKat), CNVkit (https://github.com/etal/cnvkit), R/ComplexHeatmap package (https://github.com/jokergoo/ComplexHeatmap), R/RCircos package (https://github.com/cran/RCircos), Pyclone-vi (https://github.com/Roth-Lab/pyclone-vi), EXPANDS (https://github.com/noemiandor/expands), TitanCNA (https://github.com/gavinha/TitanCNA), ggtree (https://github.com/YuLab-SMU/ggtree), Gardner-Altman plots (https://github.com/ACCLAB/dabestr), MutationalPatterns (https://github.com/UMCUGenetics/MutationalPatterns), mutationProfiles (https://github.com/nriddiford/mutationProfiles), SigneR (https://github.com/rvalieris/signeR), HDP (https://github.com/nicolaroberts/hdp), deconstructSigs (https://github.com/raerose01/deconstructSigs), sigLASSO (https://github.com/gersteinlab/siglasso), factoextra (https://github.com/kassambara/factoextra), vegan (https://github.com/vegandevs/vegan). Corrplot (https://github.com/taiyun/corrplot). The adapted code for the APOBEC3G signature is available at https://github.com/APOBEC3G.

## Acknowledgments

B.M.F was supported by the Starr Cancer Consortium grant (I12-0030). We thank Dr. Jenny Xiang (Genomics Resources Core Facility) for whole-exome sequencing and Dr. Tuo Zhang for his input on bioinformatic analysis. We thank Dr. Edwin Sandanaraj for his input on bioinformatic analysis. We thank Bing He from the Translational Pathology Core lab at Weill Cornell Medicine. We thank Dr. John Maciejowski and Alexandra Dananberg for the kind gift of the doxycycline-inducible GFP-tagged APOBEC3G vector and PiggyBac transposase vector and their input and advice on the immunofluorescence microscopy (DeltaVision Elite system). We thank Dr. Zhengming Chen for his input on statistical analyses. We thank Dr. Sushmita Mukherjee for her input and advice on confocal laser scanning microscopy, 3D reconstruction by Imaris, and the quantification of immunofluorescent images. We thank the Microscopy and Image Analysis Core Facility. Finally, we thank Dr. Nathaniel Landau for his input and advice.

## Author Contributions

Initiation and study design: W.L., K.P.N., and B.M.F.

Mouse experiments: K.P.N., L.B., S.R.R., and W.L.

Pathology analysis: F.K.

Cell experiments: D.N. and L.G.

Statistical and bioinformatic analyses: W.L., K.W.E., B.B., M.U., C.R., A.S., O.E., and B.M.F.

Analysis of HIV mutational data: L.C.

Supervision of research: B.M.F.

Writing of the first draft of the manuscript: W.L. and B.M.F.

All authors contributed to the writing and editing of the manuscript.

## Additional information

**Supplementary information** is available for this paper.

Correspondence and requests for materials should be addressed to B.M.F.

## Supplementary Table Legends

**Supplementary Table S1| Mice included in the study, survival data, and tumor stage**.

**Supplementary Table S2| Single nucleotide variants in murine bladder tumors**.

**Supplementary Table S3|96-channel trinucleotide mutational spectra for murine bladder tumors**.

**Supplementary Table S4| Primers for qPCR and genotyping**.

## Supplementary Figure Legends

**Supplementary Figure S1| APOBEC3G contributes to carcinogenesis in a murine bladder cancer model**. A) Composite bar chart representing the percentage of tumor stages in mice from the *hA3G*(+) *mA3*(−/−), *mA3*(−/−), and wild-type (C57BL/6J) groups. The non-tumor category includes benign tissue, hyperplasia, and dysplasia. B) The survival curve for the *hA3G*(+) *mA3*(−/−) (red) group, the *mA3*(−/−) (blue) group, and the wild-type (C57BL/6J) (purple) group. The *hA3G*(+) *mA3*(−/−) mice had a similar survival curve to wild-type mice. Log-rank test. All dead mice had pathologically confirmed bladder cancers. *hA3G*: human *APOBEC3G*, *mA3*: mouse *Apobec3*.

**Supplementary Figure S2| APOBEC3G is present in the nuclear compartment**. A) SDS-PAGE western blot for the murine tumor organoids derived from the *hA3G*(+) *mA3*(−/−) tumor and *mA3*(−/−) tumor. B) SDS-PAGE western blot for single-cell clones from the human urothelial bladder tumor cell line 5637 with doxycycline-inducible GFP-tagged APOBEC3G. C) SDS-PAGE western blot for single-cell clones from the human urothelial bladder cancer cell line RT112 with doxycycline-inducible GFP-tagged APOBEC3G. A3G-myc: APOBEC3G tagged with myc tag. A3G-GFP: APOBEC3G tagged with GFP. *hA3G*: human *APOBEC3G*, *mA3*: mouse *Apobec3*.

**Supplementary Figure S3| APOBEC3G is capable of entering the nuclear compartment**. A) Experimental schema for immunofluorescence imaging of extracted nuclei. B) Representative images of extracted nuclei from human urothelial bladder cancer cell line RT112 with integrated doxycycline-inducible GFP-tagged APOBEC3G vector. The extracted nuclei were stained by the DAPI and α-Tubulin as a cytoplasmic marker control. The quantitative result of the GFP signal within the nucleus is presented in a box plot as median with IQR. The lower whisker indicates Q1-1.5*IQR. The upper whisker indicates Q3+1.5*IQR. The dot indicated an individual nucleus. Multiple comparisons were performed. *: *P*<0.05. C) Experiment schema of confocal laser scanning microscopy and 3D reconstruction. D) Representative image for the 3D reconstruction of the extracted nucleus. The violet surface indicated the nuclear surface reconstructed from DAPI staining. The green signal indicated the GFP signal within the reconstructed DAPI surface. IQR: interquartile range.

**Supplementary Figure S4| APOBEC3G induces kataegic events *in vivo* identified by SeqKat**. A) Rainfall plot of kataegic loci in the *hA3G(+) mA3(−/−)* and *mA3(−/−)* tumors. The kataegic loci were identified by SeqKat. Vertical lines represent individual tumors. Grey dots represent singlet substitutions. Orange dots indicate the substitutions within the APOBEC-unrelated kataegic loci. Purple dots indicate substitutions within the APOBEC-related kataegic loci with a significant number of C>T or G>A in kataegic loci calculated by binormal test (**Methods**). B) Bar chart representing the number of kataegic loci indicating that APOBEC-related kataegic loci were enriched in *hA3G*(+) *mA3*(−/−) tumors. Fisher’s exact test. *: *P*<0.05. C) Strand asymmetry of C>T substitutions in APOBEC-related kataegis. Each line represents an APOBEC-related kataegic locus in different genotypes. The length of the line indicates the relative distance between substitutions. The triangles with different directions and colors indicated the strandedness of C>T substitutions. D) C>T substitutions occurring *in cis* were enriched in *hA3G*(+) *mA3*(−/−) tumors. Bar chart representing the number of C>T substitutions occurring *in cis* and the C>T substitutions occurring *in trans* or singlet within kataegic loci. Fisher’s exact test. *: *P*<0.05. *hA3G*: human *APOBEC3G*, *mA3*: mouse *Apobec3*.

**Supplementary Figure S5| Heatmap of copy number variants in bladder cancers from the *hA3G*(+) *mA3*(−/−) and *mA3*(−/−) groups**. Each tumor is indicated as one vertical lane. The heat scale indicates the corrected copy number values for each segment. Chr: Chromosome. *hA3G*: human *APOBEC3G*, *mA3*: mouse *Apobec3*. CNV: copy number variant.

**Supplementary Figure S6| Circos plots of CNVs in bladder cancers from the *hA3G*(+) *mA3*(−/−) and *mA3*(−/−) groups**. Concentric circles represent individual tumors. CNV gains are represented in red and CNV losses are represented in blue. *hA3G*: human *APOBEC3G*, *mA3*: mouse *Apobec3*. CNV: copy number variant.

**Supplementary Figure S7| APOBEC3G increases intra-tumoral clonal heterogeneity**. The figure represents analyses from the EXPANDS tool. A) The *hA3G*(+) *mA3*(−/−) tumors harbor a higher number of clones compared to the *mA3*(−/−) tumors. Mann-Whitney U test. *: *P*<0.05. Boxplots show the median and IQR. The lower whisker indicates Q1-1.5*IQR. The upper whisker indicates Q3+1.5*IQR. Individual dots indicate individual tumors. B) The *hA3G*(+) *mA3*(−/−) tumors displayed higher Shannon entropy compared to the *mA3*(−/−) tumors. Mann-Whitney U test. *: *P*<0.05. Boxplots show the median and IQR. The lower whisker indicates Q1-1.5*IQR. The upper whisker indicates Q3+1.5*IQR. Individual dots indicate individual tumors. C) The median difference in Hill number order between *hA3G*(+) *mA3*(−/−) and *mA3*(−/−) tumors are shown by the Gardner-Altman estimation plot. The mean difference is plotted as a bootstrap sampling distribution and depicted as a dot with a 90% confidence interval indicated by the ends of the vertical error bar. D) Phylogenetic trees for representative *hA3G*(+) *mA3*(−/−) and *mA3*(−/−) tumors with comparable total mutational burdens but divergent clonal evolution patterns. ‘WT’ represents the inferred wild-type genome. Branch length corresponds to the proportion of the number of shared variants. The length of the branches in different tumors is normalized to the same scale. *hA3G*: human *APOBEC3G*, *mA3*: mouse *Apobec3*. IQR: interquartile range.

**Supplementary Figure S8| APOBEC3G generates a distinct *in vivo* mutational signature**. A) The percentages of trinucleotide motifs harboring C>T substitutions to total substitutions were compared between tumors in the *hA3G*(+) *mA3*(−/−) and *mA3*(−/−) groups were compared using the Mann-Whitney U test. *: *P*<0.05. The light red background highlights the five trinucleotides’ motifs, which were significantly increased, and the light blue background highlight the motif, which was significantly decreased in tumors in the *hA3G*(+) *mA3*(−/−) group compared to the *mA3*(−/−) group. The light grey highlights motifs with no significant differences between the two genotypes. Boxplots show the median and IQR. The lower whisker indicates Q1-1.5*IQR. The upper whisker indicates Q3+1.5*IQR. B) Strand asymmetry of APOBEC3G introduced C>T substitutions. The ratio was calculated by dividing the number of C>T substitutions in the leading or the transcribed strand by the number of C>T substitutions in the lagging or untranscribed strand. Then, the ratio of *hA3G*(+) *mA3*(−/−) tumors was normalized to the mean ratio of *mA3*(−/−) tumors. The dot indicates the median normalized ratio of transcriptional and replicative strand asymmetry in *hA3G*(+) *mA3*(−/−) tumors. The line presented the IQR. C) The single base substitution signature induced by transgenic expression of human APOBEC3G (SBS.A3G), transgenic expression of human APOBEC3A (SBS.A3A), and wild-type mouse APOBEC3 (SBS.mA3). D) SBS.A3G has a low cosine similarity to SBS.A3A and SBS.mA3. E) Cosine similarity of experimentally derived and COSMIC PCAWG single base substitution signatures. SBS.A3G has a low cosine similarity to SBS2 and SBS13. SBS.A3A had a high cosine similarity with SBS2. *hA3G*: human *APOBEC3G*, *mA3*: mouse *Apobec3*, SBS: single base substitution, COSMIC: Catalogue Of Somatic Mutations in Cancer, PCAWG: the Pan-Cancer Analysis of Whole Genomes. IQR: interquartile range.

**Supplementary Figure S9| C>T substitutions in the *pol* gene of HIV**. The number of C>T substitutions in the *pol* gene of HIV.

**Supplementary Figure S10| Validation of signature induced by APOBEC3G**. A) The single base substitution signature induced by human APOBEC3G (SBS.A3G) was extracted using the statistical framework based on the mutational shift distance (MSD), hierarchical Dirichlet process (HDP), and Bayesian treated non-negative matrix factorization (NMF) model (SigneR) (**Methods**). B) SBS.A3G extracted using the three different tools have high cosine similarity. C) Cosine similarity of the signatures extracted from experimental data with known COSMIC PCAWG single base substitution signatures. *hA3G*: human *APOBEC3G*, *mA3*: mouse *Apobec3*, SBS: single base substitution, COSMIC: Catalogue Of Somatic Mutations in Cancer, PCAWG: the Pan-Cancer Analysis of Whole Genomes. IQR: interquartile range.

**Supplementary Figure S11| *APOBEC3A*, *APOBEC3B*, *APOBEC3G* mRNA expression in CCLE cancer cell lines**. A) *APOBEC3G* mRNA expression level in different cancer types. B) *APOBEC3A*, *APOBEC3B*, and *APOBEC3G* mRNA expression in tumor cell lines. Data was obtained from CCLE. A3A: APOBEC3A, A3B: APOBEC3B, A3G: APOBEC3G.

**Supplementary Figure S12| The correlation between APOBEC3G mRNA and SBS.A3G mutational loads in TCGA pan-cancer cohorts**. A nonparametric Spearman correlation analysis was performed between *APOBEC3G* mRNA expression and SBS.A3G mutational loads in TCGA pan-cancer cohorts. Each dot represents a cancer type in the pan-cancer TCGA cohort. Linear regression line (red line) and 95% CI (grey) were generated. CI: confidence interval.

**Supplementary Figure S13| The SBS.A3G signature contributes to the mutational load in human cancers**. A) The area of each circle represents the proportion of tumors with SBS.A3G contribution for each cancer type. The circle’s color represents the median count of SBS.A3G mutations in each cancer type. MSD: a statistical framework based on the mutational shift distance to extract mutational signatures. HDP: a framework to extract the signature based on the hierarchical Dirichlet process. SigneR: a framework to extract the signature based on the Bayesian treated non-negative matrix factorization (NMF) model. DS: deconstructSigs package. MP: MutationalPatterns package. SL: sigLASSO package. TMB: Tumor mutational burden. Abbreviations for the TCGA cancer types are available at https://gdc.cancer.gov/resources-tcga-users/tcga-code-tables/tcga-study-abbreviations.

**Supplementary Figure S14| SBS.A3G contributes to BLCA patients’ overall survival**. Overall survival analysis for the BLCA cohort shows that the patients with high C>T substitutions in the CCC+CCT+TCC motifs had lower survival than patients with C>T substitutions in TCW motifs. Patients were grouped based on the ratio of C>T counts in the CCC+CCT+TCC motif and TCW motifs (cutoff is 1) into two groups based on the total mutational burden in the TCGA BLCA cohort.

